# High-speed device synchronization in optical microscopy with an open-source hardware control platform

**DOI:** 10.1101/533349

**Authors:** Marshall J. Colville, Sangwoo Park, Warren R. Zipfel, Matthew J. Paszek

## Abstract

Recent advances in fluorescence microscopy have enabled the visualization of subcellular structures at unprecedented resolution. However, the complexity of state-of-the-art microscopes has increased considerably, often requiring the precise control and synchronization of multiple peripheral devices at high speeds. Drawing inspiration from open-source prototyping systems, like the Arduino, we describe the development of a new control platform that adopts the best features of these systems – affordability, facile programmability, and flexible connectivity – but with the scientific-grade inputs and outputs (I/O) and built-in routines that are necessary to control peripherals in advanced microscopy applications. Notably, our platform includes waveform generators and I/O for point-and azimuthal-scanning of excitation in laser-based applications. As a proof of concept, we show how the integration of waveform generation, multiplexed analog outputs, and native hardware triggers into a single central hub provides a versatile platform for performing fast circle-scanning acquisitions, including ring scanning-angle interference microscopy (SAIM), total internal reflection fluorescence (TIRF and ring TIRF) microscopy, and multiangle TIRF (MA-TIRF). We also demonstrate how the low communication latency of our hardware platform can reduce image intensity and reconstruction artifacts arising from synchronization errors produced by software control. Our complete platform, including hardware design files, firmware, API, software, and all associated source code, will be hosted for community-based development and collaboration.

## Introduction

Modern imaging applications typically require the cooperative action of several independent devices, such as stages, filters, excitation modulators, and many others. Control of peripheral devices directly from the instrument computer requires independent connections, communication protocols and manufacturer-provided drivers and application programming interfaces (APIs). Integration of several peripherals in this way is technically challenging, and while software packages exist from both commercial and open-source^1^ suppliers, they each have a limited set of supported devices. Moreover, the latency introduced by serial communications, the operating system (OS), application overhead and suboptimal API implementations can have a significant effect on synchronization accuracy. Because of these limitations, the time taken to update the experimental parameters across multiple peripherals can be much greater than the individual devices’ response times, limiting the effective acquisition speed and increasing the risk of synchronization errors that could cause artifacts in the collected data.

To overcome these challenges, embedded hardware controllers centralize device control within a single unit, typically using digital triggers or analog voltages to step peripheral devices through a series of predefined states. Hardware controllers can decrease communications overhead and implementation complexity by eliminating serial communications during an experiment. For devices that support digital triggering or analog control, the update time is reduced to the controller’s response time, eliminating the additional delay of serial data transfer and decoding. A controller with sufficient speed and memory can have the entire acquisition sequence pre-programmed, allowing autonomous operation from the instrument computer, eliminating communication overhead and processing time delays at runtime.

Hobby-grade microcontroller breakout boards, such as the Arduino devices^2^, have armed individuals and communities with a flexible platform to prototype and create interactive hobbyist-and enthusiast-grade electronic devices. However, these control platforms lack the appropriate I/O and optimizations to reliably control many of the high-precision, scientific-grade peripherals in state-of-the-art microscopes. Furthermore, hobby-grade controllers usually require the development of additional hardware, application-specific firmware, and computer-side software to handle communications and programming for scientific applications. General-purpose analog output and field programmable gate array (FPGA) cards are popular alternatives that offer more powerful general-purpose I/O and system integration. However, these devices share many of the same limitations as the hobby-grade controllers. At a significantly higher cost, FPGAs are often unable to source the required voltage range or power for scientific peripherals without additional amplification^3,4^. Moreover, they still suffer the drawbacks of implementation complexity at hardware and software levels. A common limitation of each of these hardware control solutions is that while the underlying controller may be general purpose, it is implemented and developed in a single, highly specialized application. Because of this the completed systems generally lack portability, limiting transferability and community support.

Here we describe an open-source instrument-control platform that is designed with flexibility, ease of adoption, performance, and cost in mind. Throughout the development of our device, we have committed to 3 major design principles: 1) the device should be highly hardware independent and support a wide range of peripherals, 2) the device should not limit data acquisition rate and 3) the device should be an open-source community resource. While the instrument controller is general purpose by design, specific I/O and routines have been included to provide a low-cost, high-performance solution for rapid peripheral device synchronization in advanced microscopes, including laser-scanning systems. We demonstrate the advantages of hardware control in a custom azimuthal beam scanning microscope by characterization of the effects of excitation quality and timing accuracy in a pair of complementary axial localization techniques: scanning angle interference microscopy (SAIM)^5,6^ and multiangle total internal reflection fluorescence microscopy (MA-TIRF)^7^. As our platform is open-source and built with widely available and open-source tools wherever possible, any researcher can recreate, customize, or create derivatives of our controller for community-based development.

## Results

### Controller design

The instrument controller that we developed formed a hardware abstraction layer between the computer and peripheral devices. We selected a full-speed USB HID protocol that is supported on all major operating systems (OS) for communications between the instrument computer and controller. Use of the HID protocol eliminated the need for OS-specific USB drivers, making the controller “plug and play”. While the HID protocol could provide up to a 64 kB/s bandwidth for high-speed interrupt transfers, this was implemented as 64-byte transfers sent at 1 ms intervals. With camera acquisition synchronization signals on the order of the HID polling rate, the HID protocol did not scale well with system complexity and speed. To overcome this limitation, the instrument controller used a 16-bit PIC(r) microcontroller with 256 kB program memory and 96 kB RAM in our design. The microcontroller had a sufficiently large program memory space to perform complex run-time operations and room for the addition of new features as needed. The 96 kB of RAM allowed for storage of complex, multi-parameter experimental sequences, overcoming HID bandwidth limitations. While designing the controller, we identified a set of key input/output (I/O) types that are common to a variety of instruments, including waveform generation, analog voltages, and digital I/O, and made these available on a set of generic connectors. Thus, the controller could interface with a broad range of peripheral devices (Fig. 1a, Supplementary Note 1).

**Fig. 1:**
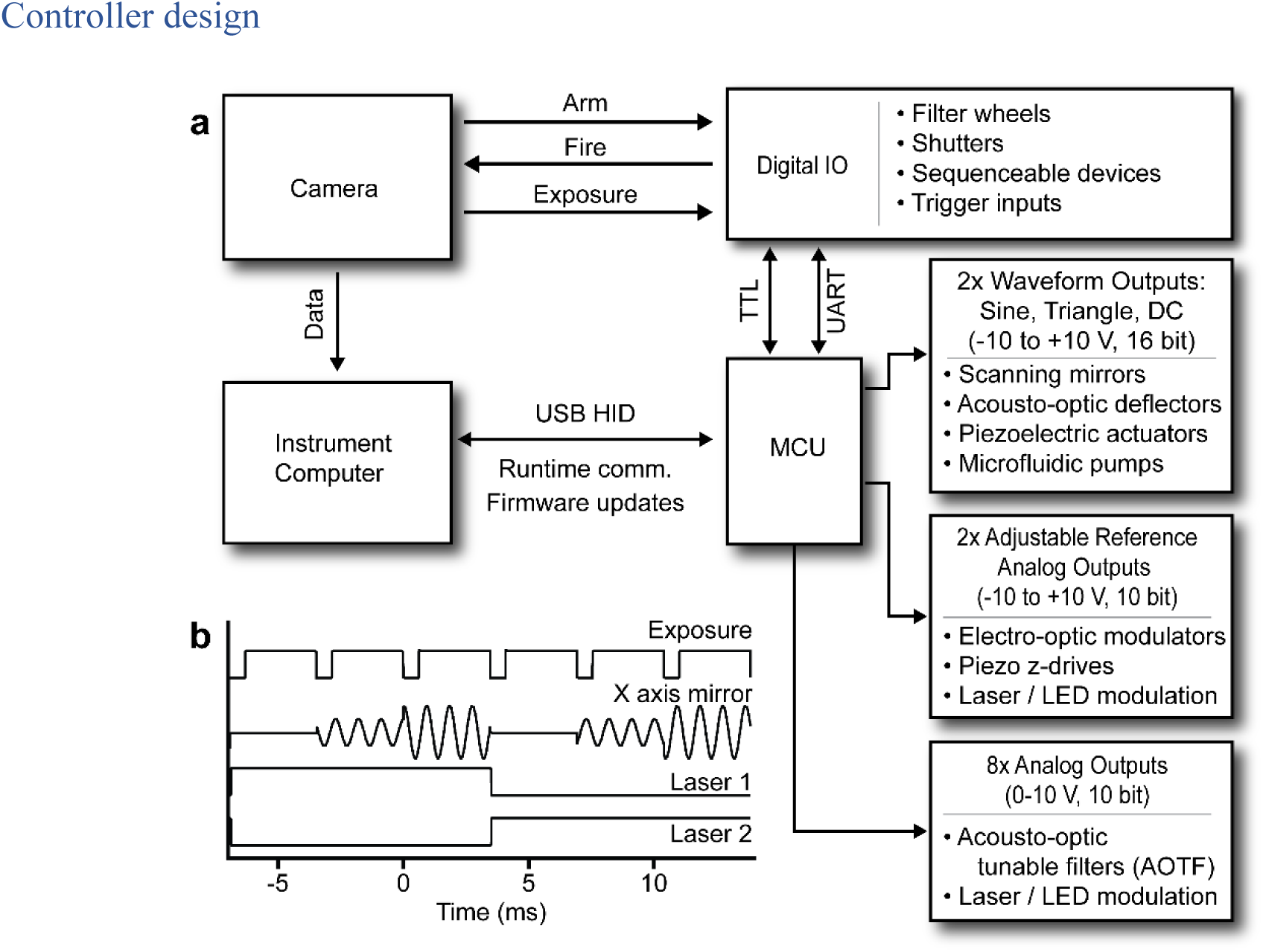
Hardware control integration facilitates the cooperation of a variety of peripheral devices. **a** Conceptual system topology wherein the micro-controller acts as an intermediary between the instrument computer and multiple peripherals. **b** Timing waveforms from a maximum framerate acquisition sequence. The camera’s exposure acts as the timing signal to synchronize peripheral device updates in a typical circle-scanning experiment. The experiment was defined as a series of 3 repeating scan radii (mirror waveform amplitude) for each of 2 excitation wavelengths (Lasers 1 and 2).

We built our hardware controller around a microcontroller to take advantage of its low cost, field reprogrammability, native interrupts, and chip-level integration of relevant functions, such as those for serial communications, counters, and parallel ports. While the simplicity of the “system-on-a-chip” design of the microcontroller was one of its strengths, it came at the expense of relatively slow computation speed. To address this, we offloaded calculations for acquisition sequence to the instrument computer. Prior to data acquisition the experimental parameters were uploaded to the controller, which ran as a finite-state machine during the experiment. The controller stepped through the experimental sequence using the microcontroller’s native interrupts triggered by the camera exposure, device triggers or internal timers (Fig. 1b). Furthermore, we prioritized the allocation of parallel ports and internal serial communication hardware to minimize processor overhead wherever possible (Supplementary Fig. 1).

We found that for a typical experimental step, including waveform amplitude change, excitation shuttering and modulating 2 lasers, the total update time was on the order of 50 μs. For reference, the vertical shift time of our electron-multiplying CCD camera (EMCCD; Andor iXon 888 Ultra) was approximately 600 μs and the readout time of our scientific CMOS (sCMOS; Andor Zyla 4.2) was approximately 3.9 ms (Supplementary Fig. 2). Thus, the controller could synchronize several peripheral devices far faster than the dead time between exposures on a state-of-the-art microscope and was capable of handling complex, experimental sequences without limiting speed.

**Fig. 2:**
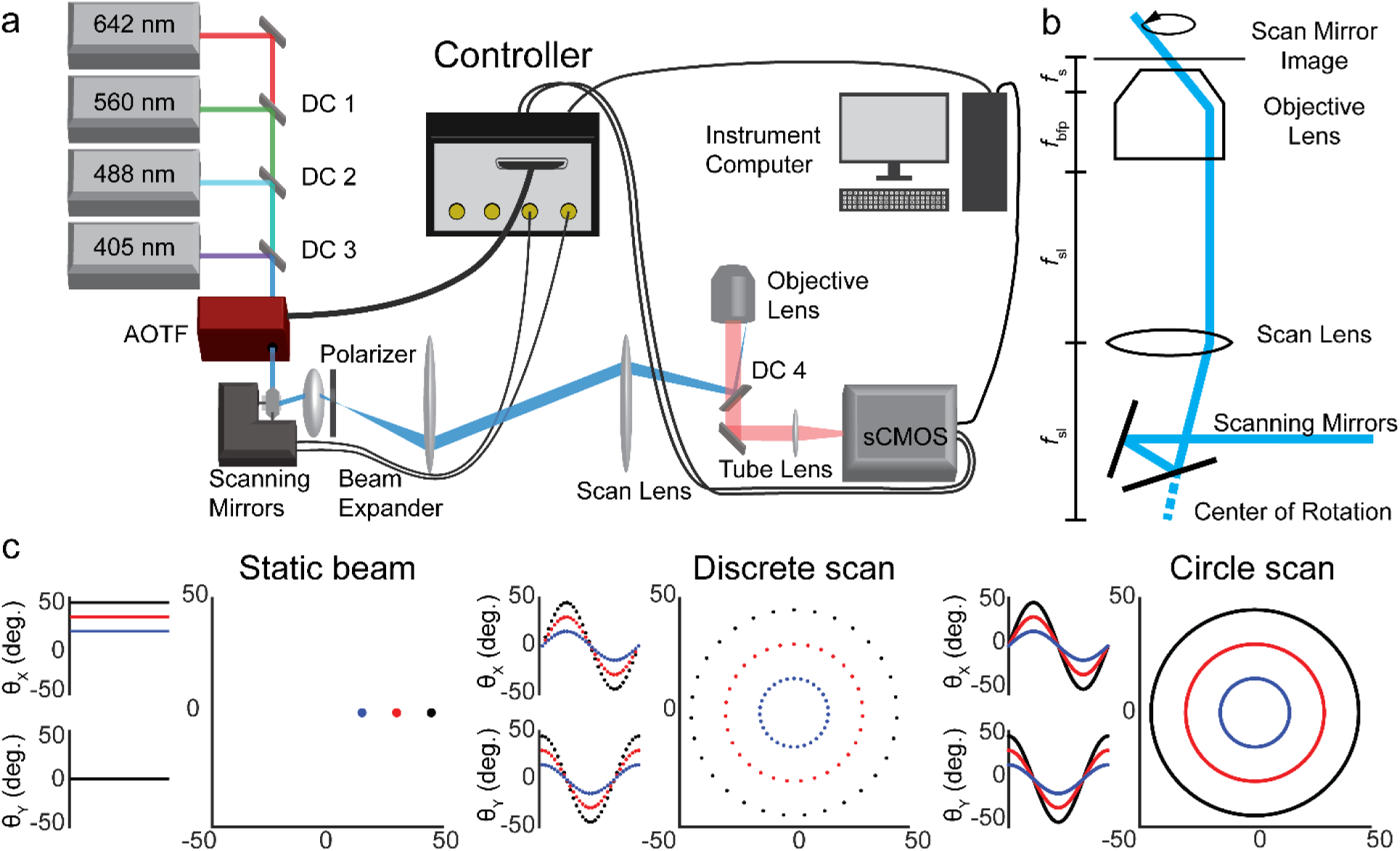
Circle scanning microscope design with hardware control. **a** Schematic illustration of the key components of the circle scanning microscope. DC1-DC3: Laser combining dichroic mirrors. AOTF: Acousto-optic tunable filter. Polarizer: m = 1 vortex half-wave retarder. DC4: Quad bandpass dichroic. Additional components omitted for clarity. **b** Conceptual diagram of the circle scanning optics. *f*sl: Scan lens focal length. *f*bfp: Objective lens rear focal length. *f*s: Objective lens front focal length. **c** Scan mirror command signals and laser pattern on the objective rear focal plane. Static beam: the laser parked at a single location on the objective back focal plane by applying a constant voltage to the galvanometer-mounted mirrors. Discrete scan: the mirrors drive the beam through 32 points approximating a circle in the objective back focal plane. Circle scan: the mirrors are driven with sinusoidal voltages scanning the beam in a continuous circle.

### Azimuthal beam scanning

Speckles and fringes caused by imperfections in the optical train, such as dust particles or internal reflections, are inherent to laser illumination^8^. The uneven excitation profile caused by these coherence artifacts is particularly noticeable in TIRF excitation and other widefield imaging techniques. While a variety of technologies have been developed to reduce or eliminate these artifacts^9–11^, azimuthal beam scanning, or “circle scanning,” has become an attractive solution to reduce the effects of interference fringes and speckle artifacts in fluorescence microscopy^12,13^. Therefore, we incorporated several laser-scanning routines into our controller and demonstrated how a super-resolution azimuthal beam scanning microscope could be built around our open-source control platform (Fig. 2a).

In our microscope, the controller formed a central hub, synchronizing the cooperative actions of the laser-scanning mirrors, acousto-optic tunable filter (AOTF), sCMOS camera and other devices (mechanical shutter, EMCCD etc., Fig. 2a). The scanning mirrors’ angular deflections were controlled by the dual waveform outputs of our controller. The scanning mirrors’ apparent center of rotation was positioned in the objective’s conjugate image plane so that the excitation beam location remained static in the sample plane regardless of the azimuthal or axial angle (Fig. 2b). This configuration of the optical system produced a homogeneous excitation field by scanning many azimuthal angles within a single camera frame, with the effective profile being the average across the azimuthal angles in each exposure. The sCMOS fire-all and EMCCD exposure outputs were both connected to one of the controller’s digital I/O ports which was configured as an external interrupt for acquisition synchronization. Using the cameras’ exposure signal allowed precise shuttering by the AOTF, only exciting the sample during the sensor active time. Additionally, peripheral updates, such as changing the scan radius or excitation wavelength, were begun immediately at the end of an exposure, ensuring the system had enough settling time before the beginning of the next exposure without adding additional delays between frames (Supplementary Fig. 2, Supplementary Note 1).

In an azimuthal scanning microscope, the excitation laser must be steered about the optical axis such that its position describes a circle in the objective lens’ back focal plane. This can be accomplished by a variety of beam steering methods including crossed acousto-optic deflectors^14^, deformable micromirror devices^7^, and motorized mirrors^13,15^. We noted that circular scans could be approximated with a discrete set of points (discrete scan, Fig. 2c and Supplementary Fig. 3a, Supplementary Note 2) lying along a circle. However, for mechanical beam steering mirrors with finite maximum accelerations, we found that our galvanometer-driven mirrors were unable to follow the command signal perfectly. When we used a 32-point circle approximation, the mirror momentum resulted in oscillations about each point, decreasing the precision with which the axial incidence angle at the sample could be controlled (Supplementary Fig. 3c).

**Fig. 3:**
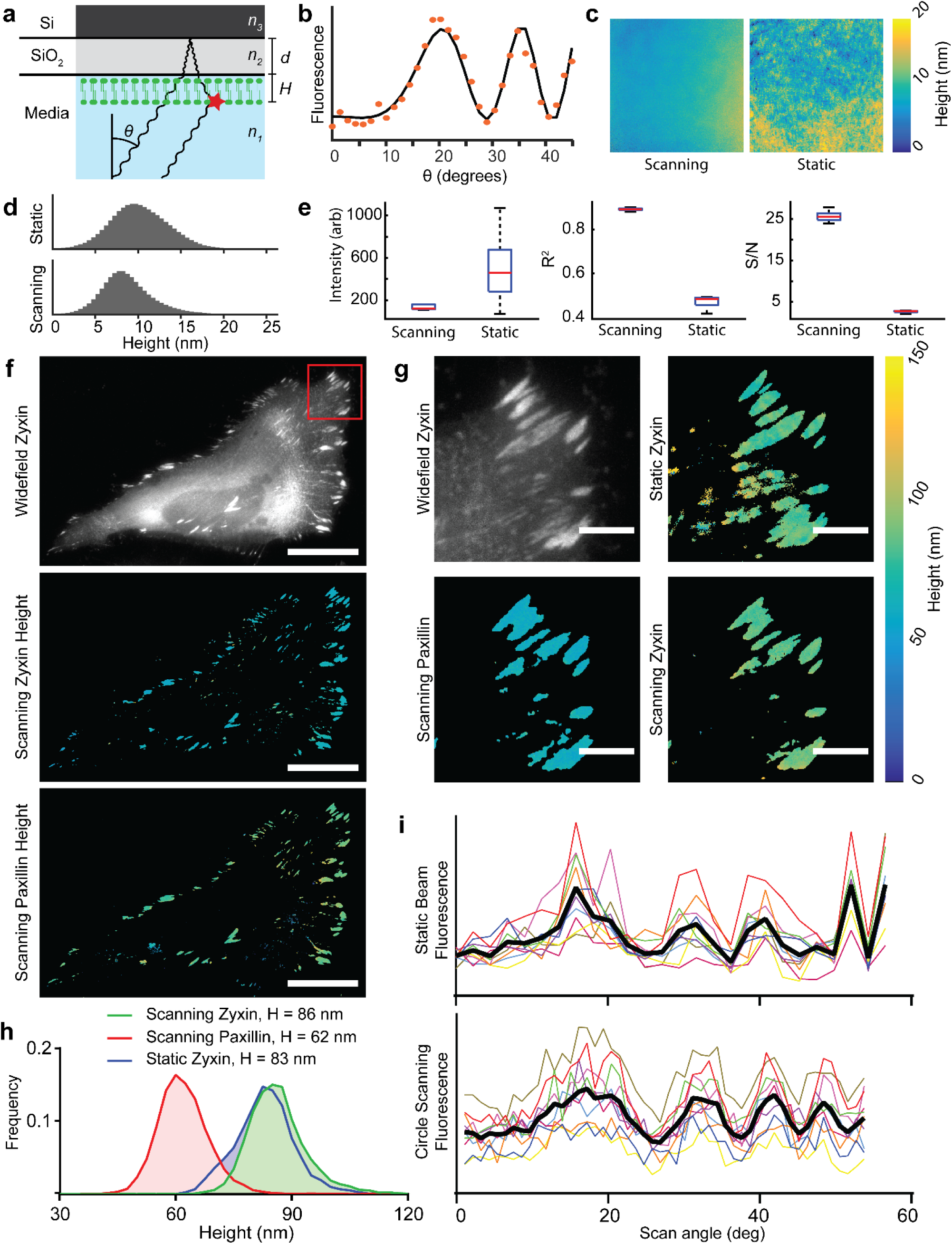
Azimuthal beam scanning reduces incidence angle dependent laser artifacts in scanning angle interference microscopy (SAIM) imaging. **a** Schematic illustration of SAIM. Samples were prepared on reflective silicon substrates with a defined oxide layer of thickness *d* and excited with a laser at incidence angle *θ* to generate an axial interference pattern with known spatial frequency. The measured fluorescence intensity at distance from the substrate *H* varies with *θ* for a supported lipid bilayer (SLB) labeled with DiI. **b** Example single-pixel raw data (orange dots) and best fit curve. **c** Reconstructed SLB height maps with circle scanning or static beam excitation schemes (full image 147.9 x 147.9 μm). **d** Pooled DiI height distributions in SLBs acquired with static beam and circle scanning excitation schemes (n = 5 regions, 4.2*10^6^ pixels per region). **e** Quantification of the average fit parameters in circle scanning and static beam SAIM imaging of the 5 lipid bilayer regions in **d**. Red lines indicate mean, boxes the central quartiles, whiskers min and max. **f** Widefield and circle scanning SAIM height reconstructions of a live HeLa cell expressing the focal adhesion components mEmerald-Zyxin and mCherry-Paxillin, scale bars 25 μm. **g** Enlarged view of the region boxed in **f**, scale bars 5 μm. A static beam reconstruction of zyxin height is included for comparison. **h** Distribution of pixelwise height fits from adhesion complexes in **c**. Heights are the average of all non-zero fits. **i** Plots of fluorescence intensity as a function of scan angle for the static and circle scanning Zyxin images in **g**. Colored lines indicate the same 10 representative pixels from each image, black lines the average of all 10 pixels.

As an alternative strategy, we used the integrated waveform generators of our control platform to drive the mirrors with a pair of sine waves with a π/2 phase offset (circle scan, Fig. 2c and Supplementary Fig. 3b). With this approach the mirrors remained settled on the circle and in constant motion, which eliminated the noise associated with starting and stopping at each point in a discreet scan (Supplementary Fig. 3c,d).

Utilizing the integrated waveform generators in circle scanning mode, we investigated the improvements that could be gained through circle scanning in SAIM experiments to minimize laser-induced interference artifacts. SAIM is a surface-generated interference localization microscopy technique that measures the pixelwise average axial position of fluorescent probes with better than 10 nm precision. SAIM samples were prepared on a reflective silicon wafer with a defined layer of thermal oxide to act as a transparent spacer (Fig. 3a). When illuminated with a coherent light source, the direct and reflected excitation created an axial interference pattern with a characteristic spatial frequency that depended on the axial angle of incidence^16^. A series of images at known axial incidence angles were acquired and fit pixelwise to an optical model that predicted the observed fluorescence intensity fluctuations as a function of axial position (height) above the SiO2/water interface (Fig. 3b). Previous SAIM implementations utilized a static beam excitation scheme^5^, the simplest method to implement on a commercial microscope.

We hypothesized that circle scanning SAIM would remove the pixelwise, frame-to-frame variations in the excitation intensity arising from speckle and fringe artifacts in a static beam system that degrade the accuracy of axial position reconstructions. To illustrate the advantages of circle scanning over static beam excitation in SAIM experiments, we used our microscope to image supported lipid bilayers (SLBs) labeled with DiI using either a static beam or circle scanning excitation scheme. The bilayer topography from static beam reconstructions showed local topographical variations of several nanometers which were not present in circle scanning reconstructions (Fig. 3c). When multiple, independent regions of the samples were imaged, we found that the same topographical patterns emerged, indicating that the excitation artifacts caused erroneous height estimations in the analysis (Supplementary Fig. 4a). Figure 3d shows the height distributions from the combined results of 5 bilayer regions with average heights of ± 3.46 nm and 8.72 ± 3.10 nm for the static beam and circle scanning experiments, respectively. To verify that the samples were equivalent in terms of labeling density and bilayer continuity we examined the fluorescence intensity maps (Supplementary Fig. 4b). Based on a visual inspection, the height reconstructions in both cases were independent of variations in label density or bilayer defects. The average of 5 height reconstructions showed the topographical variations seen in the static beam experiments were consistent across multiple regions unlike the circle scanning experiments, indicating that they were systematic and did not reflect the sample topography (Supplementary Fig. 4c). This led us to conclude that the topography observed in the static beam experiments was a consequence of laser excitation artifacts which are mitigated by circle scanning. The improvement in the quality of the raw data was reflected in the more consistent average intensity across image sets and higher average R^2^ value from the reconstructions (Fig. 3e). Finally, circle scanning resulted in a greater than 5-fold increase in the average signal to noise ratio (S/N) across the experiments, a direct consequence of eliminating the incidence angle dependent laser artifacts.

**Fig. 4:**
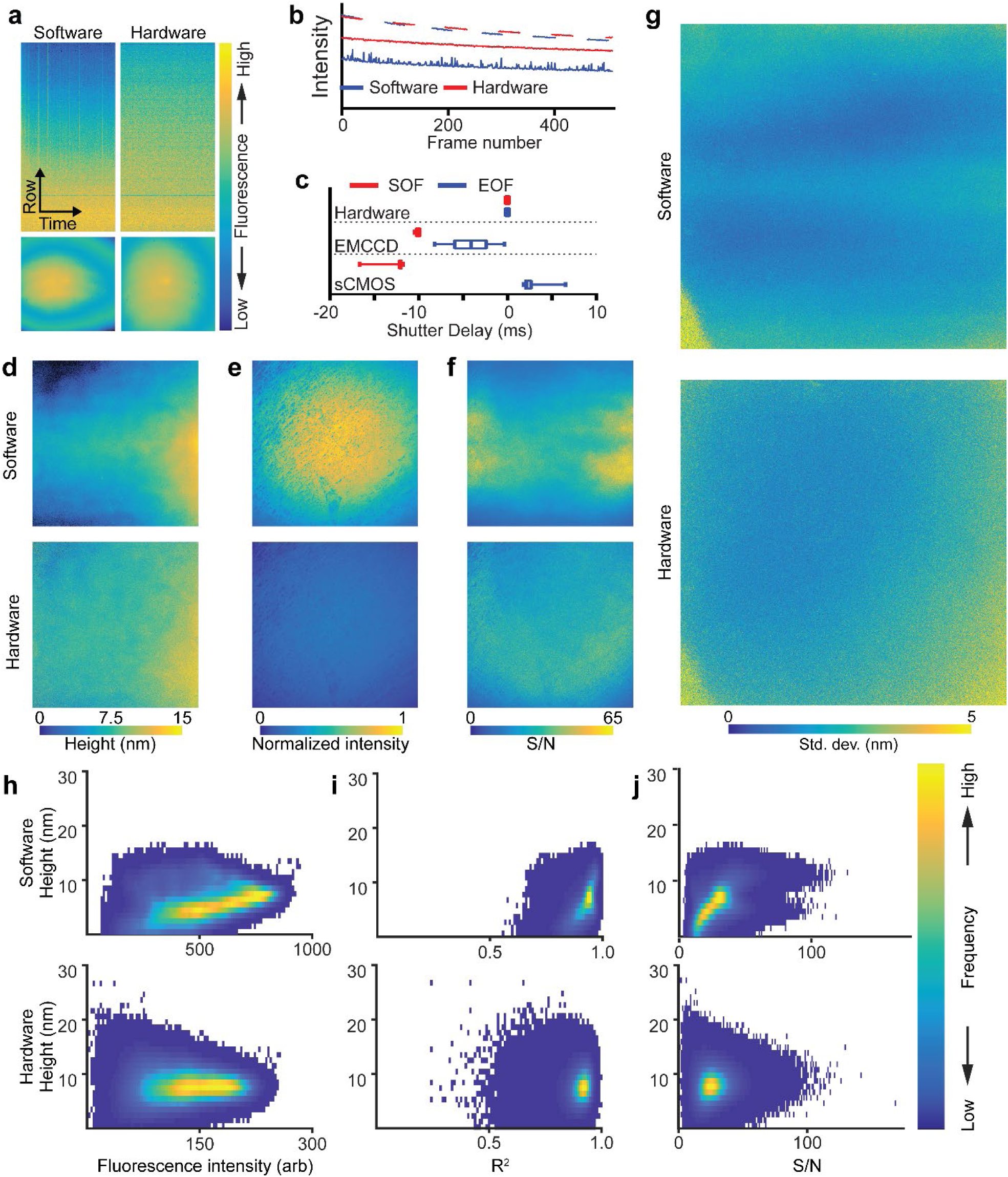
Hardware based synchronization minimizes latency-induced artifacts in SAIM. **a** Top: fluorescence kymographs of the top half of the center pixel column of an sCMOS camera imaging a homogeneous dye monolayer. Bottom: Full sensor single frame images from the same acquisition sequence. **b** Plot of average row intensities (2048 pixels) from the series in **a**. Dashed lines represent the center pixel row, solid lines the top pixel row. **c** Distribution of excitation shutter delays at the start of frame (SOF) and end of frame (EOF) relative to the camera’s exposure signal over 50 frames. Lines represent the average of 50 frames, boxes the center quartiles and whiskers the minimum and maximum values. **d-f** Lipid bilayer height, fluorescence intensity and signal-to-noise ratio SAIM reconstructions from image sets acquired with software or hardware synchronization. **g** Standard deviation from SAIM reconstructions of lipid bilayer height. Each of 6 equivalent regions of the sample were imaged first with hardware synchronization and then immediately after with software synchronization. The horizontal bands in the software image follow the center-out readout pattern of the sCMOS camera. **h-j** Lipid bilayer height distributions in SAIM reconstructions vs. fluorescence intensity, coefficient of determination and signal-to-noise ratio with the excitation under software (top row) or hardware (bottom row) control. Images in **a** and **d-g** reflect the full sensor area of 2048 x 2048 pixels, corresponding to a 147.9 x 147.9 μm field of view. Data in **a** and **b** were normalized to the average intensity in the first exposure for comparison.

When imaging biological samples, such as live cells, high background fluorescence, local variations of the sample’s optical properties caused by intracellular structures and dynamic processes conspire to disrupt the ideal axial interference pattern. While these properties cannot be eliminated as they are intrinsic to the sample, optimizing instrument performance could possibly mitigate their effects. To demonstrate the advantages of circle scanning in a biologically relevant context, we imaged live HeLa cells expressing the focal adhesion proteins mEmerald-zyxin and mCherry-paxillin (Fig. 3f). We noted that paxillin and zyxin could serve as axial measuring sticks as their height distributions are well known from several sources^17–19^. The widefield image in Fig. 3f highlights the background fluorescence in a typical live-cell sample. Because SAIM illuminated throughout the sample volume, the fluorescence from diffuse pools of free probes within the sample contributed to the observed intensity at each incidence angle. For bright, discrete structures, such as focal adhesions on the cell periphery, the intensity fluctuations of the background were typically much smaller than the signal magnitude and robust localization was possible. With dim structures or in regions with no structure, fluctuations in the background, such as those arising from laser excitation artifacts, could be erroneously identified as real structures in the reconstruction. In the magnified view of the cell’s leading edge (Fig. 3g), the widefield image showed several adhesion complexes on the periphery of the cell and a typical inhomogeneous background away from the cell’s edge. The laser artifacts in the static beam case resulted in the artifactual appearance of structures that were not visible in the widefield image. The circle scanning reconstructions were free of these errors and matched the focal adhesions from the widefield image. Both circle scanning and static beam reconstructions of zyxin showed similar distributions in height of 86.4 ± 8.17 nm and 83.1 ± 7.69 nm, respectively, as expected (Fig. 3h). Inspecting the pixelwise raw data demonstrated the improved S/N and reduction in outlier data points at high incidence angles where laser illumination artifacts are particularly problematic (Fig. 3i).

### Accuracy of peripheral device synchronization in hardware versus software control

Another potential source of error in SAIM and other techniques that make frame-to-frame intensity comparisons challenging is variation in the excitation dose. Excitation shuttering minimizes photobleaching and phototoxicity in living samples, reducing the total light dose to the minimum required for the experiment. However, the excitation shutter must be accurately synchronized with the camera exposure. Variations in shutter/camera synchronization between frames are reflected in the observed fluorescence intensity, leading to localization artifacts. This effect is particularly problematic when using sCMOS cameras with rolling shutters where the beginning of exposure and readout are done by pixel rows^20^. Shuttering partway through the exposure/readout cycle results in the over exposure of some rows, while exposure throughout the exposure/readout cycle can lead to spatial distortion of fast-moving objects. sCMOS cameras can be run in a simulated global shutter mode wherein the excitation exposure is begun once all pixel rows are exposing and ended when the first row begins to readout (i.e. “fire-all” mode).

We considered that the roundtrip communication travel time from camera to computer to AOTF and the variable latency inherent in software control schemes could limit the precision of shutter synchronization. We collected 500 frames of a homogeneous fluorescent monolayer with either software control or hardware control to illustrate the effects of synchronization errors with a sCMOS camera (Fig. 4a). From the pixel column kymograph, with software control the upper pixel rows were relatively underexposed compared to the center pixel row. Under hardware control this effect was reduced, and a more even fluorescence intensity pattern was observed. Timing variability had the greatest effect on the last rows to be exposed or readout (those furthest from the center of the sensor in a center-out readout configuration, Fig. 4b). The center row (dashed lines in Fig. 4b) showed little difference between software and hardware control schemes while the top row (solid lines) had a lower average value and greatly increased noise under software control when compared to hardware control (Fig. 4a,b). To quantify the synchronization errors, we placed a fast photomultiplier tube directly in the excitation path of our microscope (Fig. 4c, Supplementary Fig. 5). With software synchronization, we measured an average −12.34 ± 0.9943 ms and 2.714 ± 1.236 ms delay between the beginning or end of exposure, respectively, and shutter operation. On the other hand, when the camera’s fire-all signal was used to trigger the excitation shutter through our hardware controller, we achieved a 3 order of magnitude improvement in synchronization latency (12.73 ± 12.80 μs and 10.59 ± 1.552 μs start and end of exposure, respectively, Fig. 4c, Fig. S5). With careful calibration the average delay under software control could be reduced. However, the variability due to communications and software overhead were intrinsic and could not be easily accounted for. Our controller utilized a hardware interrupt, enabling or disabling the shutter in just a few processor clock cycles to achieve very precise timing while being immune to the complexities that limit software synchronization accuracy.

**Fig. 5:**
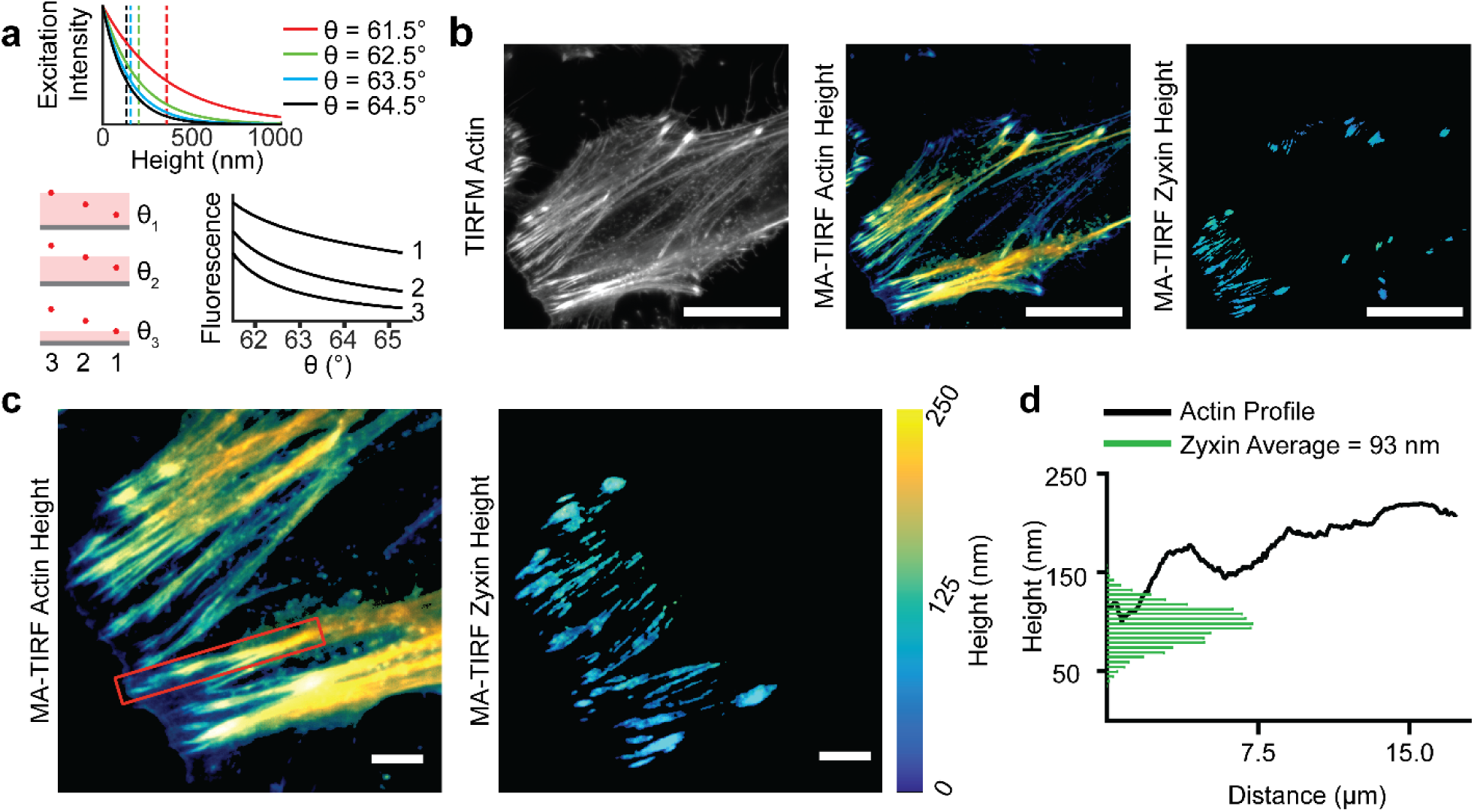
Implementation of circle scanning multiangle TIRF (MA-TIRF). **a** Concept of MA-TIRF. Top: The evanescent excitation field intensity (solid lines) as a function of height above the glass-liquid interface and penetration depth (dashed lines) for various angles of incidence. Lower left: Schematic representation of fluorescent probes (red dots) at 300, 200 and 100 nm (3, 2 and 1, respectively) from the glass-liquid interface. Lower right: Theoretical intensity profiles as a function of angle of incidence for the 3 fluorescent probes. **b** Widefield and height reconstructions of phalloidin-AF560 and mEmerald-Zyxin in a fixed HeLa cell. Scale bars 25 μm. **c** Enlarged view of the leading edge of the cell in **b**. Scale bars 5 μm. **d** Reconstructed actin height profile (solid black line, red box in **c**) and zyxin height distribution (green bars, non-zero fits from **c**).

Considering these results, we hypothesized that the increased precision of hardware synchronization would result in a higher S/N and better localization precision in SAIM. To make a direct comparison between software and hardware synchronization we imaged SLBs first with software synchronization and then, without moving the sample, with hardware synchronization at the same excitation intensity and over the same range of angles, such that an identical observation of the sample was made. The height reconstructions of SLBs (Fig. 4d) acquired with software synchronization demonstrated a strong dependence of fitted height on distance from the center of the sensor which was not reflected in the hardware synchronization acquisitions. The fitted fluorescence intensity profile (Fig. 4e) was closely matched with both synchronization schemes. The hardware acquisition sequences had a lower average intensity, which was expected given the shorter overall excitation time (Fig. 4c). The S/N from the reconstructions with software synchronization showed a high S/N band in the region around the center of the detector that decays toward the top and bottom of the image compared to the hardware synchronization acquisition where the S/N closely followed the excitation intensity profile, as expected (Fig. 4f). Using the standard deviation in height over several independent SLB regions as a metric for localization precision, we found that the sensor readout pattern was reflected in the localization precision with software synchronization and not with hardware synchronization (Fig. 4g). Owing to the large sensor size in the sCMOS camera, the excitation profile was not perfectly flat across the field of view, which could make interpretation of the reconstructions difficult. To better understand the effects of synchronization errors, we examined the height distributions as a function of intensity, coefficient of determination (R^2^, a measure of goodness of fit) and S/N (Fig. 3h-j). In the case of software synchronization, the reconstructed heights showed a strong dependence on the fluorescence intensity with higher intensity corresponding to larger height values. A similar trend was observed for both R^2^ and S/N. With hardware synchronization the reported height was independent of intensity and similar distributions were seen for both R^2^ and S/N.

### MA-TIRF

To demonstrate some of the versatility of our hardware control platform, we performed MA-TIRF imaging experiments utilizing the same hardware and sequencing functions used in SAIM experiments. Like SAIM, MA-TIRF achieves axial localization of fluorescent probes by reconstruction from multiple independent images. When the excitation incidence angle at the coverslip to sample interface is beyond the critical angle, θc, total internal reflection occurs, and the excitation potential is confined to a thin evanescent field within a few hundred nanometers of the interface. The exponential decay of the evanescent field depends on the angle of incidence and is characterized by the penetration depth, which decreases with angle of incidence^21^. In an MA-TIRF experiment the excitation incidence is swept through a range of angles greater than the critical angle and individual images are acquired at each step (Fig. 5 a).

As a test subject, we imaged actin stress fibers, which are involved in adhesion-based motility and are required for maintenance of focal adhesion complexes. The ubiquity of actin fibers throughout cells complicates methods such as SAIM which lack optical sectioning. The thin evanescent excitation field in MA-TIRF rendered the ventral stress fibers clearly visible. We also imaged zyxin, which is known to interact with actin in the upper layers of focal adhesions where the mechanical coupling between the cytoskeleton and adhesion machinery occurs. Simultaneous imaging of actin and zyxin highlighted MA-TIRF’s ability to resolve stress fiber profiles (Fig. 5 b, c). Compared to SAIM, MA-TIRF returned a similar average zyxin height in adhesion complexes (93.3 nm) but a broader standard deviation (19.9 nm vs. 8.17 for circle scanning SAIM). Using MA-TIRF we were able to visualize the height profile along actin stress fibers (Fig. 5 d) originating in adhesion zones as determined by zyxin enrichment. We found that in the zyxin-rich regions the stress fiber height matches the average zyxin height but increased with distance from the zyxin end. The ability to perform both SAIM and MA-TIRF experiments on the same microscope platform demonstrated the ability of our hardware controller to quickly adapt existing infrastructure for new applications.

## Discussion

High-speed synchronization of multiple devices has become a common necessity in the field of microscopy and presents a significant barrier to entry for many researchers needing to build or customize prototype microscopes. The two common approaches to peripheral integration are through software packages or dedicated hardware controllers. Software is typically the simplest path to system integration, but generally limits the available hardware pool to a list of supported devices. In some cases, such as the open-source Micro-Manager^1^ microscopy software, unsupported devices can be added by writing new device adapters. Doing so requires familiarity with the specific programming languages used, the device and software APIs and in some cases the operating system architecture. Furthermore, inefficient or improper implementations can limit performance and affect data quality. As we have shown in Fig. 4, even in the case of a stable and widely used software package, limitations imposed by the communication protocol’s speed and intricacies can limit the synchronization precision of multiple devices.

Hardware device synchronization can overcome many of the limitations imposed by software. We demonstrated this for the example case of camera and AOTF timing wherein the excitation is shuttered in concert with the camera’s actual exposure. A naïve solution would be to avoid shuttering the excitation at the cost of increased excitation dose, photobleaching and phototoxicity. The greater the interframe interval the more pronounced each of these effects becomes, making it infeasible for time-course experiments with significant delays between frames. Under software control and with shuttering we observed timing differences of 10 or more milliseconds between the exposure and shuttering. If synchronization differences are consistent they can be accounted for in software by adding a delay between the slow and fast devices. In the case we have presented, latency between camera and AOTF events had variabilities on the order of milliseconds, and a simple delay would not suffice. Under hardware control we were able to achieve precise and consistent synchronization regardless of the framerate

We have used lipid bilayers to demonstrate that small variances in the timing between excitation and exposure can decrease the accuracy and precision of SAIM experiments. SLBs formed by fusion of small unilamellar vesicles make an ideal test sample with known properties (e.g. thickness, index of refraction, structure), are easily labeled with a variety of dies and have extremely low background fluorescence. The SLB experiments illustrate the importance of frame-to-frame consistency in maximizing the precision of SAIM localizations. Inconsistent illumination caused by interference at the sample (laser fringes and speckles) has the same effect as inconsistent timing on SAIM experiments. Although the causes are different, the result is that on a pixelwise basis, the instrument is causing additional intensity variations that obscure the predicted intensity variations in the raw data. In this study we chose not to use a set of background or correction images in any experiment. We found that the combined circle scanning and hardware synchronization approach eliminated the need for preprocessing the data with correction images.

Resolution of fast biological processes places additional constraints on the imaging system, and the effective time resolution is limited to the elapsed time to acquire the complete image series for a range of scan angles. Ultimately, this limit is set by the camera framerate plus any delay associated with changing the excitation angle or wavelength. In a typical circle scanning SAIM experiment we acquire between 16 and 32 images over the incidence angle series. The exact number of images is determined empirically at the time of the experiment based on balancing acquisition time with data quality. Our hardware controller allows acquisition at the camera framerate, that is there is no additional delay added and all devices are update during readout. Under these conditions the effective time resolution is the number of frames per acquisition multiplied by the exposure period. For instruments with slower peripheral devices (i.e. motorized TIRF prisms), or under software control, an adequate delay between frames must be added while the system reaches the new stable point for the next camera frame. Motion of the sample between the first and last frames of a SAIM or MA-TIRF sequence will introduce additional errors in the reconstruction because a pixel in the first frame is not mapped to the same sample location in subsequent frames in the angle series. It follows that the time between the first exposure and last exposure in a set of incidence angles should be less than the time it takes the sample to move by one pixel. This condition can be difficult to satisfy but hardware control maximizes the achievable acquisition rate, providing a more accurate multi-frame snapshot in SAIM and MA-TIRF experiments (Supplementary Fig. 6, Supplementary Videos 2,3).

There are many forms of hardware control that have been used in microscopy from simple hobby-grade microcontroller platforms to commercial field programmable gate array boards to custom embedded electronics platforms covering a broad range of cost, complexity and flexibility. We have sought to create a broadly useful platform that is economical and easy to integrate. The open source model used by Micro-Manager has greatly contributed to the success of the project and we seek to emulate their example by making all design and source code freely available^22,23^. We have attempted to use freely available and open-source tools wherever possible in the development of the project to encourage adoption by other researchers. We have made all levels of the project from hardware to driver API and user interface software open-source. This will allow us to capitalize on the skills and feedback of users and create a community of contributors, distributing the cost of further development based on the framework presented here. Furthermore, while we have demonstrated the utility of our controller in a single, circle scanning instrument the control platform we have developed has a broad range of applications including laser-scanning confocal or two photon microscopies, imaging cytometry, microfluidic systems and spectroscopy to name a few. Through continuing, community-driven development of the project we envision our hardware as a valuable resource for general scientific device control.

## Methods

### 1. Preparation of supported lipid bilayers

A chloroform solution of 1-palmitoyl-2-oleoyl-glycero-3-phosphocholine with 0.1 mole percent 1,2-dipalmitoyl-sn-glycero-3-phosphoethanolamine-N-(cap biotinyl) (Avanti Polar Lipids) was dried first under a stream of nitrogen and then under vacuum to remove the solvent. The resulting film was dissolved to 1 mg/mL in PBS pH 7.4 at 37 °C with extensive vortexing and freeze-thawed 7x by cycling between liquid nitrogen and 37 °C baths. The lipid suspension was then extruded 13x through double stacked 100 nm polycarbonate membranes (Whatman) to yield small unilamellar vesicles (SUVs). SUVs prepared in this way were stored at 4 °C and used within 2 weeks. Silicon wafers with ∼1900 nm thermal oxide (Addison Engineering) were diced into 1 cm^2^ chips, and the oxide layer thickness for each chip was measured with a FilMetrics F50-EXR. The chips were then cleaned in piranha solution (30 volume % hydrogen peroxide in sulfuric acid), rinsed extensively and stored in deionized water until use. SLBs were formed by incubating the cleaned silicon chips in the SUV solution for 1 hour, and then rinsed with PBS, incubated with 1.75 μM DiIC18 for 30 minutes, and rinsed again with PBS. The samples were then inverted into a 35 mm glass-bottom imaging dish (MatTek) containing PBS and immediately imaged.

### 2. Preparation of HeLa cell samples

The plasmids mEmerald-Zyxin-6 (Addgene plasmid # 54319; http://n2t.net/addgene:54319; RRID: Addgene_54319) and mCherry-Paxillin-22 (Addgene plasmid # 55114; http://n2t.net/addgene:55114; RRID: Addgene_55114) were a gift from Michael Davidson. HeLa cells (ATCC CCL-2) were cultured in Dulbecco’s Modified Eagle Medium supplemented with 10% fetal bovine serum and 100 U/mL penicillin-streptomycin (ThermoFisher) at 37 °C and 5% CO2.

For SAIM experiments, silicon chips were prepared as for SLBs except the chips were sonicated for 15 mins in acetone, 15 minutes in 1 M NaOH and UV sterilized in place of the piranha cleaning. Cells were seeded onto the chips at 2×10^4^ cm^-2^ in full culture medium and incubated overnight. Cells were transfected with 1:1 mEmerald-Zyxin-6 to mCherry-Paxillin-22 using Lipofectamine 3000 (ThermoFisher) per the manufacturer’s protocol and allowed to grow for 24 hours. Sample were then rinsed 3x in PBS, inverted into a 35 mm glass-bottom imaging dish containing HEPES-Tyrode’s (119 mM NaCl, 5 mM KCl, 25 mM HEPES buffer, 2 mM CaCl_2_, 2 mM MgCl_2_, 6 g/L glucose, pH 7.4) and imaged at 37 °C.

For MA-TIRF experiments, cells were seeded at 2×10^4^ cm^-2^ directly into 35 mm glass-bottom imaging dishes and incubated overnight to allow attachment. Samples were then transfected with mEmerald-Zyxin-6 as before and incubated an additional 24 hours. Following incubation, the samples were rinsed with PBS and fixed in 4% formaldehyde in PBS for 10 minutes, rinsed 3 additional times and permeabilized with 0.1% v/v Triton X-100 in PBS for 10 minutes. The solution was then aspirated, and the sample was incubated for 1 hour at room temperature with Alexa Fluor(tm) 568 Phalloidin (ThermoFisher) diluted to 1 U/mL in PBS with 1% bovine serum albumin (ThermoFisher). Samples were then rinsed 3 times in PBS and imaged immediately.

### 3. Controller Design

Full details on the controller design are included in Supplementary Note 1 and 3. Briefly, the analog subcircuits were designed and simulated using CircuitLab^24^. The controller PCB layout was made using EAGLE (Autodesk). Printed circuit boards were ordered unpopulated from OSH Park. The components were purchased from Digi-Key and then hand assembled. Firmware was written in the C programming language, compiled and debugged with C-Aware IDE (Custom Computer Service Inc.). Host-side driver and GUI software was written in C++ with Visual Studio 2017 (Microsoft) using the Qt^25^ framework, HIDAPI^26^, boost C++^27^, and OpenCV libraries^28^.

### 4. Microscope Design

The circle scanning microscope was built on a Nikon Ti-E inverted fluorescence microscope. The 405 nm (Power Technology Inc.), 488 nm (Coherent), and 560 nm (APB) lasers were combined with the 642 nm (APB) to make a colinear beam by a series of dichroic mirrors (Chroma). The combined output beam was attenuated and shuttered by an AOTF (AA Opto-Electronic) before being directed onto the galvanometer scanning mirrors (Cambridge Technology). The image of the laser on the scanning mirrors was magnified and relayed to the sample by 2 4*f* lens systems, the beam expanding telescope and the scan lens / objective lens combination. The beam expander is formed by *f* = 30 mm and *f* = 300 mm achromatic lenses with a m = 1 zero-order vortex half-wave plate positioned between them and positioned 2*f* from the 300 mm achromatic scan lens (Thorlabs). SAIM experiments were performed with a 60x N.A. 1.27 water immersion objective, and MA-TIRF with a 60x N.A. 1.49 oil immersion objective lens (Nikon). Fluorescence emission was collected with a quad band filter cube and single band filters (TRF89901-EMv2, Chroma) mounted in a motorized filter wheel (Sutter). For both SAIM and MA-TIRF experiments, images were acquired with a Zyla 4.2 sCMOS camera using the microscope’s 1.5x magnifier for a total magnification of 90x. The camera, filter wheel and AOTF (software control) were controlled through Micro-Manager^1^. Details on excitation incidence angle calibration are given in Supplementary Note 4.

### 5. SAIM

For static beam SAIM experiments, a series of 31 images were acquired at evenly spaced incidence angles from −45.8 to 43.9 degrees. For circle scanning experiments, 32 images were acquired at evenly spaced intervals from 0 to 30 degrees in the case of live cells or 0 to 45 degrees in the case of SLBs. Angle ranges are given with respect to the wafer normal in the imaging media (Fig. 3 c). The raw images were fit pixelwise without further preprocessing using a custom analysis program written by the authors (Supplementary Note 5) to obtain the reconstructed height topography of the samples. Images were postprocessed with Fiji^29^ and statistical analysis was performed in MATLAB (MathWorks) and Prism (GraphPad).

### 6. MA-TIRF

For MA-TIRF experiments, a series of 20 images were acquired from 61.5 degrees to 65.3 at intervals of 0.2 degrees with respect to the excitation angle of incidence in the coverslip at the glass to water interface. The galvanometer command amplitude corresponding to the critical angle was experimentally confirmed using the fluorescence intensity of a dye monolayer^30^. MA-TIRF reconstructions were done with MATLAB using the ADMM^31^. The resulting height images were postprocessed in Fiji and statistical analysis was performed in Prism.

### 7. Code availability

All source code written by the authors including controller firmware, driver, graphical interface and SAIM analysis code is available in the project repository at git@github.com:mjc449/SAIMscannerV3.git. Additional source code and third-party libraries are available from the links referenced in the controller design section. Analysis code for the MA-TRIF experiments is available at git@github.com:zcshinee/Pol-TIRF.git.

## Data availability

Source data including raw and processed image files are available upon request from the corresponding authors.

## Acknowledgements

We thank Susan Daniel and Rohit Singh (Cornell University) for discussions and assistance preparing SLB samples. This research was funded by the National Institute of General Medical Sciences Ruth L. Kirschstein National Research Service Award 2T32GM008267 (M.J.C.), National Science Foundation Graduate Research Fellowship DGE-1650441 (M.J.C.), National Cancer Institute U54 CS210184 (M.J.P.) and R33-CA193043 (M.J.P. and W.R.Z.), and the Kavli Institute at Cornell for Nanoscale Science. This work was performed in part at the Cornell NanoScale Science & Technology Facility (CNF), a member of the National Nanotechnology Coordinated Infrastructure (NNCI), which is supported by the National Science Foundation (Grant NNCI-1542081).

## Contributions

M.J.C., W.R.Z. and M.J.P. conceived the project and devised the studies. M.J.C. and W.R.Z. conceived and developed the controller electronics and firmware. M.J.C. built the controller and microscope, wrote the driver, interface and analysis software. M.J.C. and S.P. performed the experiments and collected instrument performance data. All authors contributed to the writing of the manuscript.

