## Supplementary Information and Figures for "High-speed device synchronization in optical microscopy with an open-source hardware control platform"

### Supplementary Note 1

The controller electronics are divided into analog, digital I/O, in-circuit serial programming (ICSP) and microcontroller (MCU) subcircuits. Fig. S1 provides a conceptual schematic of the analog subcircuit designs. Fig. S7 shows the populated PCB assembly highlighting the various subcircuits in the physical layout. The complete schematics and design files are available at git@github.com:mjc449/SAIMscannerV3.git. Supplementary video 1 shows a brief demonstration of high-speed beam steering and excitation synchronization.

#### Waveform generators

The analog waveform outputs consist of a pair of matched subcircuits and are designed to provide accurate synchronization while still being independently controllable. To accomplish this each has independent DC references and direct digital synthesis (DDS) components. The output frequency and phase of each DDS is set by the MCU with ranges of ~0.1 to $12.5*10^6$ Hz and 0 to 2π, respectively. Because the DDS units share a 25 MHz master clock the relative frequency and phase settings are matched between the units, preventing drift. The DDS units have independent serial data lines to the MCU so that each can be programmed individually while maintaining common frame synchronization and serial clock signals. In this way the phase, frequency and waveform type for each unit can be updated simultaneously, ensuring consistent phase values and timing. The waveform output of the DDS units has a positive bias and relatively small amplitude of ~650 mV which is insufficient to directly control most peripheral devices. To precondition the DDS output the signal amplitude is increased and a bias is applied by a precision amplifier for each channel. The amplification and bias are controlled by trimming potentiometers on the circuit board. The waveform output amplitude from the controller is proportional to the conditioned waveform reference with 16-bit resolution over the reference voltage range. The tunable amplifier circuit therefore provides a means to set a constant DC bias to the waveform reference signals as well as limit the maximum and minimum output values without decreasing the output resolution. For applications where a high-resolution DC signal is required the waveform generators can be disabled by setting the DDS units in reset (midscale output). Each of the two waveform outputs are controlled by independent dual-channel digital to analog converters (DACs). For each output's DAC one reference channel is the conditioned DDS. The second DAC channel uses a precision, trimmable 10 V reference and operates in a bipolar mode to provide a full-scale range of -10 to 10 V DC. At the output stage the two channels of each DAC are summed in a current to voltage output amplifier. The dual-channel DACs share a 16-bit parallel port on the MCU and common output register load signal. The DACs can be programmed independently or simultaneously by channels, allowing both independent and concurrent updates. Values are written to the outputs simultaneously by the common load signal, eliminating delays and ensuring output synchronization.

##### Bipolar analog outputs

In addition to the precision, 16-bit waveform analog outputs we have included a pair of 10-bit analog outputs with user adjustable references. Each channel has an independent 10 V reference and operates in bipolar mode. The references are connected to the DAC's reference inputs through a user-selectable jumper and trimming potentiometer. The jumper can be selected to bypass the center tap on the potentiometer to set the DAC output to a full-scale range of -10 to 10 V. Alternatively, the jumper can be set to connect the potentiometer center tap to the DAC channel's reference input. In this configuration the potentiometer forms a voltage divider, setting the reference value ($V_{ref}$) from ~0.1 to 10 V. The DAC output range in this mode is limited to -$V_{ref}$ to +$V_{ref}$, providing the full, 10-bit resolution across a wide spectrum of applications. The DAC is connected to one of the MUC parallel ports to maximize update speeds.

#### 8-channel 0-10 V analog outputs

The 8-channel voltage outputs are generated by pair of 4-channel DACs with a common 10 V reference. The DACs share a parallel data port from the MCU and data is written to each channel individually. However, both DACs are connected to a common output register load signal. Channels can be set individually, updating as soon as the new data is written, or simultaneously where the data for any number of the channels is loaded into the DACs then all outputs are set simultaneously. At the board level there is also a 10 V digital output and external power supply passthrough on the same pin header as the analog outputs. We use the 10 V digital line as a global shutter for our AOTF.

#### Digital I/O

We define 2 types of digital I/O for the purposes of this discussion: general purpose I/O (GPIO) and triggers. GPIO lines are those which the MCU reads based on polling the state of the pin within the thread of execution. Triggers are pins that are enabled as hardware interrupts. Interrupt events (change of pin state) trigger the MCU to halt its current thread of execution and jump to the interrupt service routine (ISR) associated with the interrupt source. When the ISR completes, the MCU continues the main thread of execution from where it left off. Our controller has 3 trigger pins that can be set programmatically to function as either triggers or GPIO and 5 additional GPIO pins. Of these pins, 4 GPIO pins and 2 triggers are +5.5 V tolerant and can operate on 3.3 V or 5 V logic without the need for logic-level conversion. There are an additional 4 digital I/O lines on an RJ45 connector. These can be set programmatically to GPIO, triggers, or utilize the MCU serial communications hardware for linking other devices, such as a second controller or other custom peripherals using SPI or UART.

##### Connectors

We have avoided the use of obscure or proprietary connectors in the design of the controller (Fig. S7). The waveform and bipolar analog outputs have SMA type connectors which can easily be adapted to BNC, SMB or many other coaxial types. The 8-channel analog outputs, digital I/O and ICSP have 0.1" pin headers at the board level for maximum versatility. For example, in our completed controller the digital I/O are broken out into 2 DB-9 connectors, 4 +5.5 V tolerant GPIO lines on one and the triggers and remaining GPIO line on another. The 8-channel analog output, 10 V digital pin and external power supply pins have been connected to a DB-25 in a pin arrangement matching our AOTF analog control connector.

#### Supplementary Note 2

Galvanometer scanning mirrors are economical and easy to integrate into optical systems such as the microscope demonstrated in this study. Commercial solutions that come packaged with appropriate driver circuitry are available from several suppliers and can even be salvaged from retired confocal and other laser scanning systems. While acousto-optic deflectors (AODs) have much faster response times, the challenge of integration of two orthogonal AODs for 2D beam scanning and higher price tag make them less attractive to most builders of circle scanning microscopes. One of the significant drawbacks of galvanometer scanning mirrors is that they are mechanical devices and as such have much slower responses to changes in position than AODs. Furthermore, the momentum of the mirrors requires that the driver electronics can quickly supply large currents proportional to the step size of the command voltage when making a large step between points such as in a discrete scan at a TIRF radius.

Discrete scans ideally should match the number and frequency of scan points to some factor of the small-step response time ($T_S$), the 99% settling time, of the galvanometers. In the case of those used in our microscope this is 120 μs. To maintain the mirrors in constant motion, the time between steps along the circle must be at least $T_S/2$, or 60 μs in our case, however faster sampling will improve the quality of the waveform scanned. Ideally the update period would be $<< T_s$.

Another consideration in constructing a circle scanning microscope is the circular frequency. If the camera frame rate is on the order of the time to complete a single azimuthal scan the circular frequency must be matched to the frame rate or the benefits of circle scanning in terms of eliminating interference in the sample plane are decreased. When they are not matched for some portion of the exposure only a part of the azimuthal scan is completed and any variation in the excitation profile will be reflected in the fluorescence image. Matching the camera framerate to the scanning frequency is technically challenging. A simple solution to this problem is to maintain a scan frequency that is much greater than the minimum framerate such that the laser beam completes multiple scans in any given exposure. In this way the partial scan at the end of exposure has a negligible effect on the excitation profile.

To keep the mirrors settled on the desired waveform, the time between changes in the command signal must be held constant or fluctuations in the update rate will be reflected in the motion of the mirrors. Considering the case of a USB HID interface with the controller or driver circuitry this is difficult to achieve, as interrupt exchanges occur at a rate of 1 kHz. If the analog voltage, as in the case of our controller, has 16-bit precision this equates to a maximum of 16 points per transfer (64 bytes per packet / (2 bytes per axis * 2 axes) at 1 kHz, or 62.5 μs per scan location. This also assumes that there are no missed transfers and that the latency in processing time of each report is less than the time between the report receipt and the previous change in the command signal. The constant timing constraint is further complicated by other tasks the controller is required to perform, and a high-priority timer interrupt is the most reliable solution to this problem. While interrupts take priority over the current computation, care must be taken in programming to ensure that the controller spends the minimum amount of time in the interrupt service routine (ISR) or other processes may incur an unacceptable delay while the ISR completes. Furthermore, a race condition exists between the priority level of scanning mirror updates and USB packet processing, as neither is more important than the other. With these constraints in place it is infeasible to construct a system wherein the controller relies on the instrument computer for continuous transfer of the scan coordinates.

To circumvent the problems associated with USB transfers the entire set of discrete scan coordinates can be preloaded into the controller before the experiment begins. In this case the controller's memory should ideally store all coordinates for the experiment, and at minimum the integer number of circles to be scanned at the reliable transfer rate. Take, for instance, a real-world scenario in which the controller can reliably receive and respond to 1 in 4 in reports, or 16 kB/s and each circle contains 32 points. In this case 2 successful transfers are required per scan radius, limiting the system to a minimum of 8 ms per exposure neglecting processing time for the new radius, system update time and settling time on the new scan waveform.

A more reliable mode of operation is to have all waveforms preprogrammed in the controller before the experiment begins. For an acquisition of 32 points per radius (as in the discrete scans presented in the main text, which is still insufficient to keep the mirrors settled on the scan waveform as demonstrated in Fig. S3) with 32 individual radii 4 kB of memory is occupied by the discrete scan points alone. In a microcontroller with limited memory the number of discrete scan points and number of scan radii quickly becomes the limiting factor as far as the experimental complexity that can be achieved.

In our controller the memory requirement is reduced to 4 bytes per scan radius when using the integrated waveform generators. The integrated direct digital synthesizers (DDS) run on a dedicated 25 MHz clock. Regardless of the system state the waveforms have a constant frequency and phase, the result of which is a constant circular scan of the laser independent of processor load. This also simplifies the process of settling the mirrors on the new scan radius during an update. By referencing the output DAC to the waveform input any changes are immediately reflected in the command signal sent to the mirrors. The result is a system in which the mirrors are settled on the new scan radius with a delay of $T_s$, which is the shortest possible for a mechanical mirror system.

#### Supplementary Note 3

Software control is a popular option for many researchers because of its ease of implementation. For many peripheral devices all that is necessary is a driver installation and/or configuring the control software. Hardware control can be significantly more difficult to accomplish. Our controller is designed to be both a development platform and a general solution for instrument control. The CAD files and bill of materials are freely available in our project repository and the populated board can be purchased fully assembled from several sources such as circuithub.com, where we have posted the necessary project files[1].

The controller firmware is available through our project repository and inexpensive programming hardware can be purchased from Microchip to flash the latest firmware into the MCU. We have also created a graphic interface that simplifies creating and executing circle scanning experiments as well as general system control (Fig. S8). The software handles all aspects of communication and control of the hardware, including calibration, alignment and synchronization.

Our controller is also a development platform for prototyping instrumentation. We have conscientiously chosen to use open-source and freely available tools wherever possible. Initially the controller firmware was written with a proprietary compiler, but we are now working on a port of the firmware code to a free, C-language compiler.

The features and applications of the controller presented in this work are only a subset of its capabilities and we envision the hardware platform being useful in many other applications. The success of many open-source projects is the collaborative input and development efforts of users and contributors. Through open-source development our project can benefit from the expertise and feedback of users in fields other than our own, expanding on the capabilities and functionality of the controller.

#### Supplementary Note 4

Both SAIM and MA-TIRF rely on accurate knowledge of the angle of incidence of the excitation beam. Calibration of the beam deflection as a function of command voltage is critical in both techniques. In the past we calibrated the system with a protractor placed on the microscope stage, measuring the beam deflection at several voltages. With the development of the graphical interface software for our controller, we built in an image-based calibration tool for circle scanning. A piece of fluorescent acrylic is placed on the microscope stage directly above the objective and the laser is scanned in the plastic. An inexpensive webcam connected to the instrument computer captures images of the laser at various waveform output amplitudes directly through the controller software. The software then finds the edges of the scanned laser in the image and calculates the effective calibration value based on the index of refraction of the acrylic and the imaging medium. For SAIM experiments the imaging medium was assumed to have an index of refraction of 1.34. For MA-TRIF the calibration was performed with a sample index of refraction of 1.52 corresponding to the index of refraction of the glass coverslip.

The range of angles used in MA-TIRF is beyond those measurable using the laser emission from the objective. Before MA-TIRF experiments, we verified the calibration by measuring the fluorescence of an absorbed monolayer of IgG-Alexa Fluor 568 (ThermoFisher). As the incidence angle approaches the critical angle the excitation intensity at the glass to water interface increases rapidly, and past the critical angle the intensity falls off[2]. The fluorescent monolayer was imaged from sub- to super-critical angles in steps of 0.1 degrees. The critical angle was determined as the angle at which the maximum fluorescence emission was observed. The experimentally determined critical angle was then used to verify the image-based calibration.

#### Supplementary Note 5

The SAIM optical model is based on the fluorescence interference-contrast microscopy and has been reported many times in the literature[3–5]. We use the simplified mathematical representation of Carbone et. Al. in a highly optimized fitting program written in C++ with the Math Kernel Library (Intel). The program uses an unbounded trust-region solver, and as such the choice of initial parameter values can lead to erroneous outputs corresponding to the nearest local minimum of the nonlinear least squares problem rather than the true value. For the samples used in this study we set the initial value of height to the average values known from the literature[6]. We have found that the solution is relatively insensitive to the initial values of intensity (A) and background (B). However, a good approximation of intensity is 80% of the difference between the highest value and lowest value in the image series for a given pixel and background is simply the lowest value. Initially we implemented the analysis routine in a multithreaded MATLAB application, however after multiple rounds of optimization analysis of a 2048 by 2048 pixel SLB image set would take between 4 and 6 hours on a 6-core Intel Core i7 5820k processor running at 3.3 GHz. With careful optimization and the low-level memory access provided by C++ we can analyze the same dataset in < 30s. The analysis code supplied in the software repository is a simple program for analyzing a single dataset, however we have written it in such a way that the processing functions are contained in a single class for portability.

In this work we define signal to noise (S/N) as the root mean square background subtracted prediction divided by the root mean square residuals, or

$$S/N = \frac{(f(\theta_i, \beta) - B)_{rms}}{\left(I_i - f(\theta_i, \beta)\right)_{rms}}$$

Where $f(\theta_i, \beta)$ is the optical model, $\theta_i$ is the vector of incidence angles, $\beta = (A, B, H)$ is the parameter vector, $B$ is the background, and $I_i$ is the observed intensity in frame $i$. Additionally we have found the combined evaluation of $R^2$ and S/N for a given pixel a better metric of the quality of the fit than $R^2$ alone.

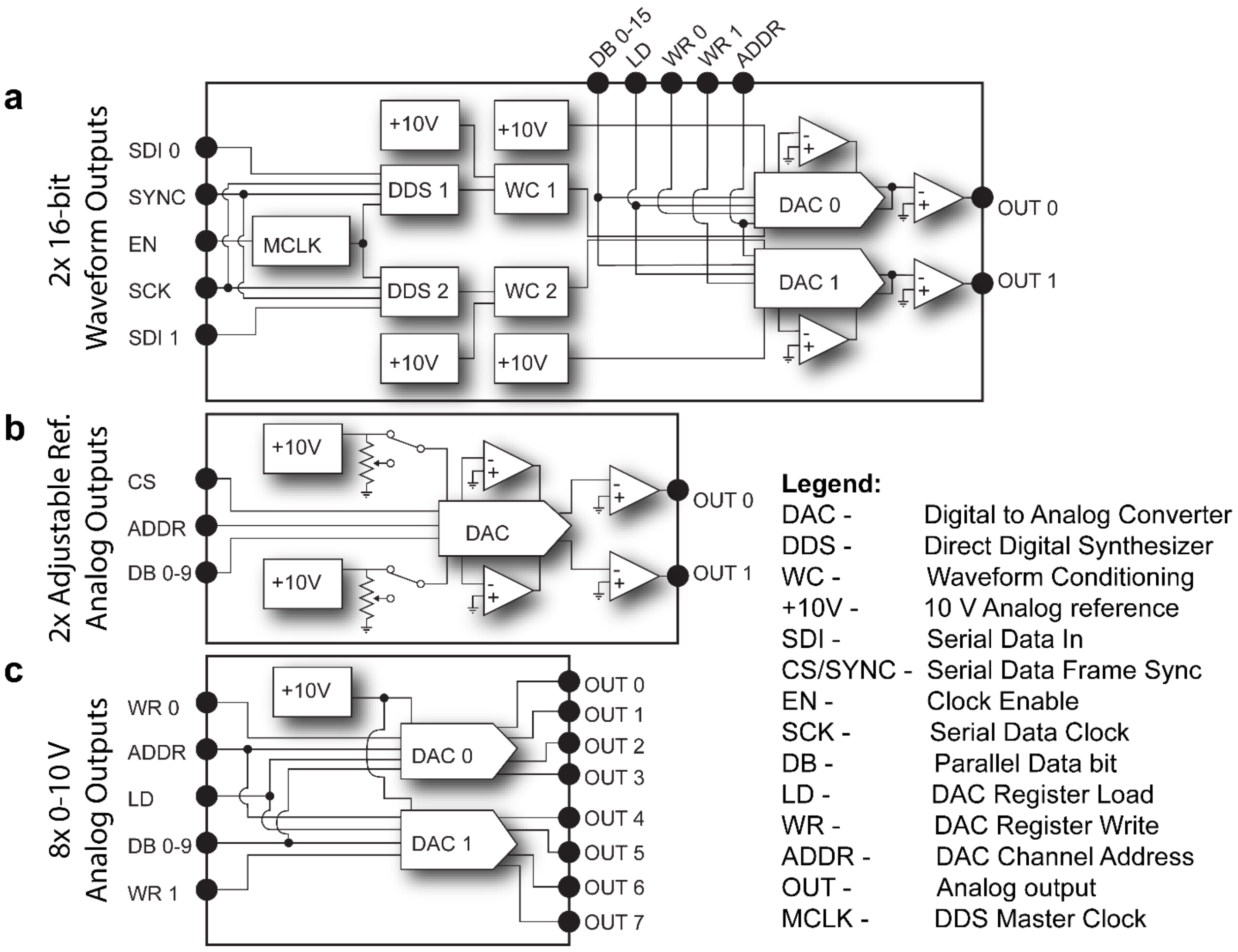


**Supplementary Fig. 1:** Expanded schematic diagrams of the controller analog subcircuits from Fig. 1. **a** The dual waveform generators share a master clock, sync and serial clock lines enabling simultaneous updates. The output amplitude and offset are individually adjusted by matched dual-channel DACs with a common load signal. The AC and DC outputs for each channel are summed in the final amplifier stage. **b** Auxiliary DAC outputs have individual, high precision 10 V references with selectable voltage dividers to tune the full-scale output range from ±0 to ±10 V. **c** Multichannel precision 0 to 10 V outputs are generated by a pair of matched 4 channel DACs sharing a data bus and load line. Each of the 8 outputs can be updated individually or all 8 simultaneously.

**a** **b**

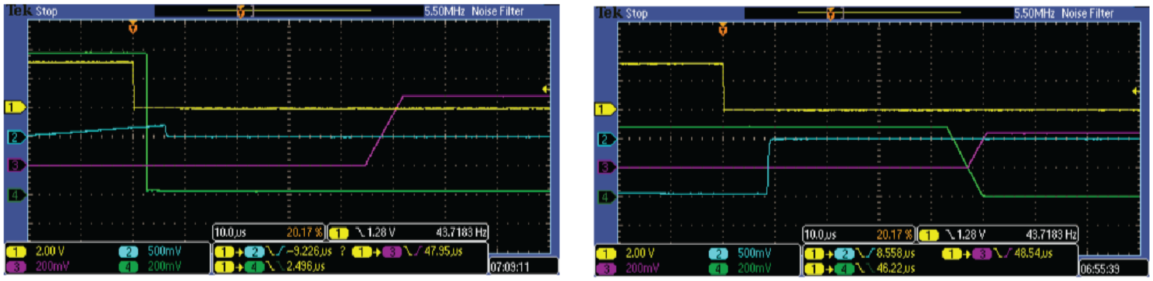


**Supplementary Fig. 2:** Example timing waveforms from a sequence acquisition. **a** The camera's exposure signal (Ch. 1, yellow) marks the end of the frame, triggering the controller to close the excitation shutter (Ch. 4, green). Immediately after shuttering the excitation the scanning mirror command waveform amplitude (Ch. 2, blue) is updated. Finally, the AOTF laser 0 channel (Ch. 3, pink) intensity is changed. The total update time is defined as the delay between the exposure signal trigger and the excitation amplitude settling on the new value. **b** Same as **a** with Ch4 (green) showing the AOTF laser 7 signal. All 8 channels of the AOTF are settled at the same time following the end of frame and the controller is ready to begin the next exposure in 50 µs.

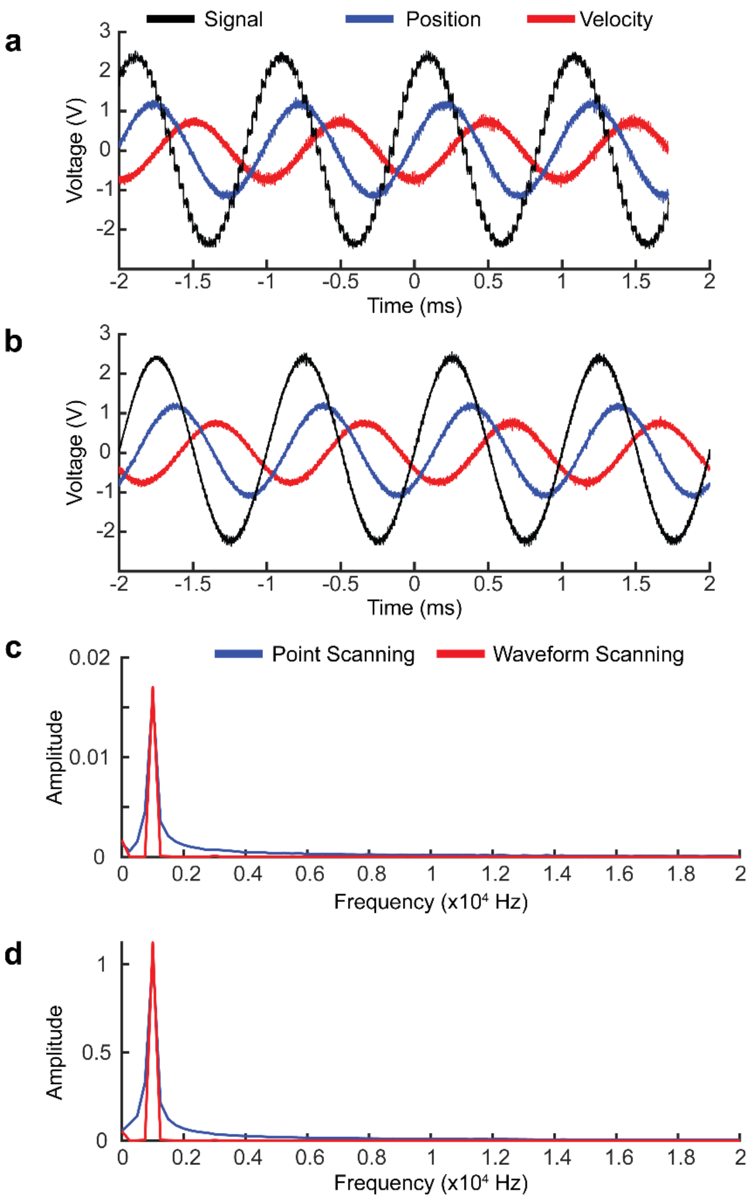


**Supplementary Fig. 3:** Galvanometer scanning mirror response to discretized (point scanning) and continuous waveforms in circle scanning experiments. **a** The scanning mirror is driven with a 32 point discretized sine wave at ~1 kHz (black trace) while it's position (blue trace) and velocity (red trace) are monitored. **b** Same as **a** when the command signal is replaced with a 1 kHz sine wave generated by the integrated waveform generators in the controller. Command signal amplitude in **a** and **b** is typical of a TIRF experiment. **c** Frequency spectra of the mirror position in a small angle scan of 1°. **d** Same as **c** with a large angle scan corresponding to a typical TIRF experiment. Increased noise in the position trace from **a** compared with **b** manifests as a broadening of the frequency spectrum in **c** and **d**.

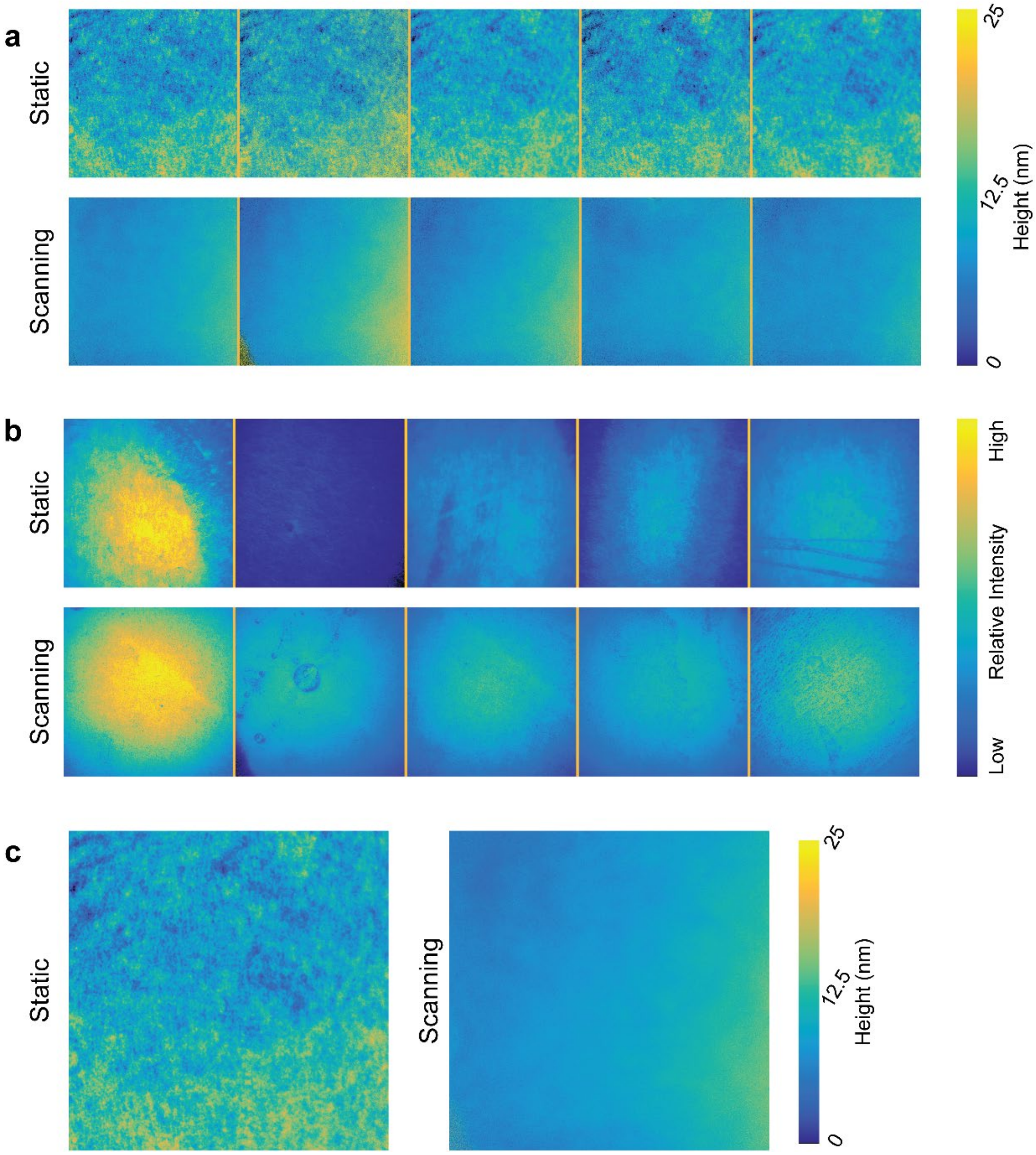


**Supplementary Fig. 4:** Laser speckle and fringing depend on incidence angle and lead to artifacts in SAIM reconstructions. **a** SAIM height reconstructions of 5 independent supported lipid bilayer regions acquired with static beam (top) or circle scanning (bottom) excitation schemes. **c** Relative intensity images (fit parameter A of the model function) of each bilayer region in **a**. Details in intensity images correspond to features in the bilayer resulting from labeling variations and bilayer defects, and are similar to a widefield image. **c** Pixelwise average height of the 5 regions in **a**. Features in the average height image are the result of pixelwise incidence angle dependent excitation variations from laser speckle and fringes. All images are the full sensor area of 2048 x 2048 pixels, corresponding to a 147.9 x 147.9 μm field of view.

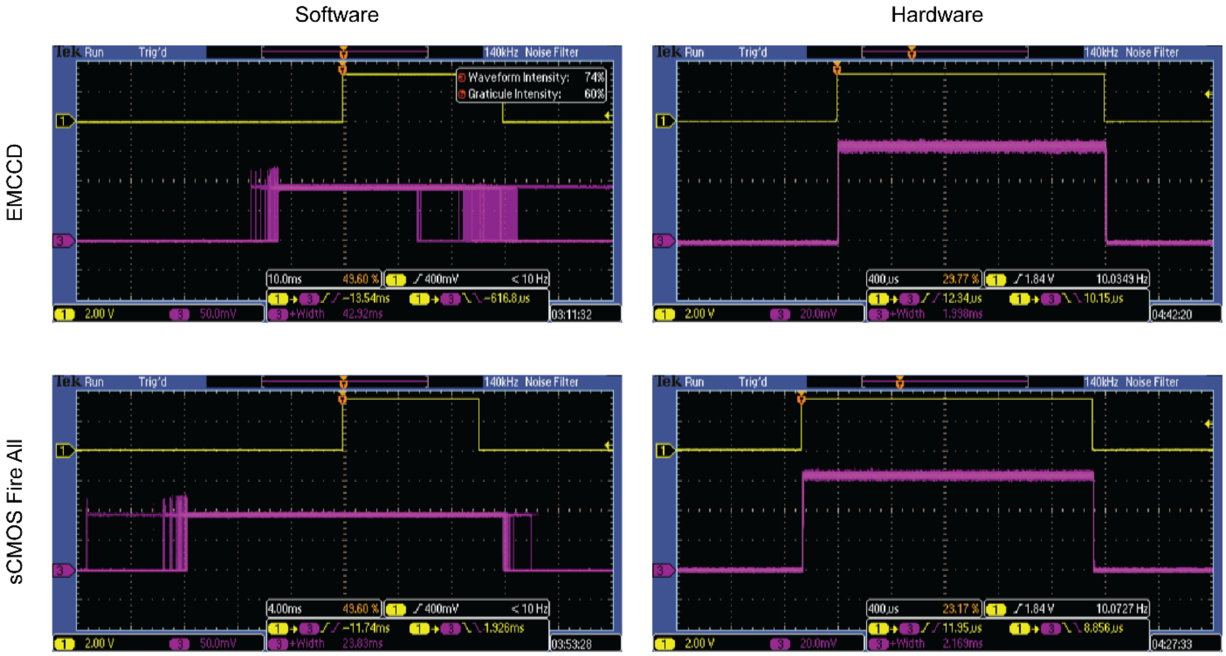


**Supplementary Fig. 5:** Excitation shuttering time comparison between software (left) and hardware (right) control. Ch1 (yellow) camera exposure signal; Ch3 (pink) excitation intensity collected by a photomultiplier tube placed directly in the excitation path. Traces from 100 individual exposures are overlaid to highlight the variability in excitation timing.

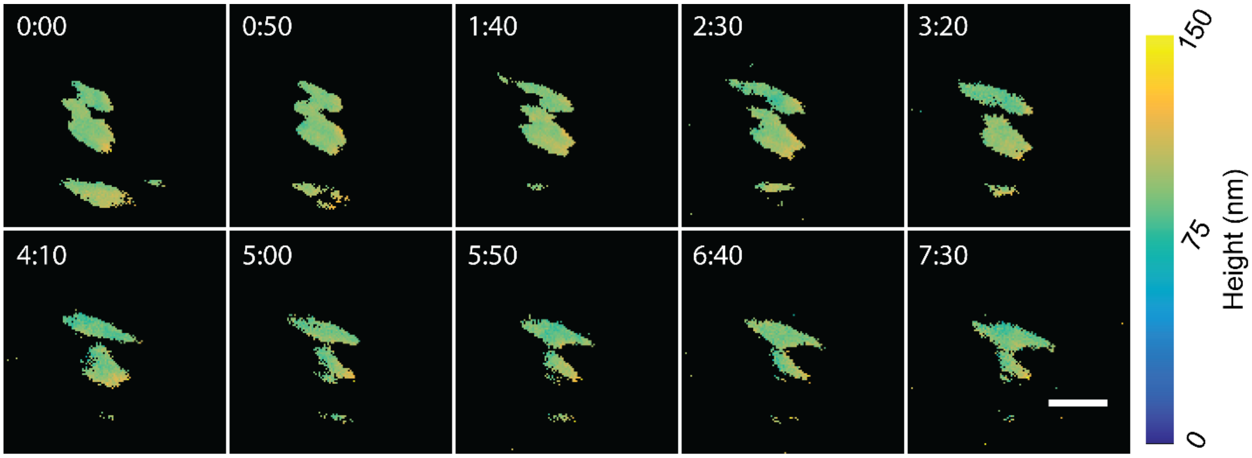


**Fig S6:** Time lapse SAIM reconstructions of the focal adhesion protein Zyxin. A live HeLa cell expressing mEmerald-Zyxin was imaged at 5 second intervals using the circle scanning excitation scheme. Scale bar 2.5 µm; time stamps are min:sec.

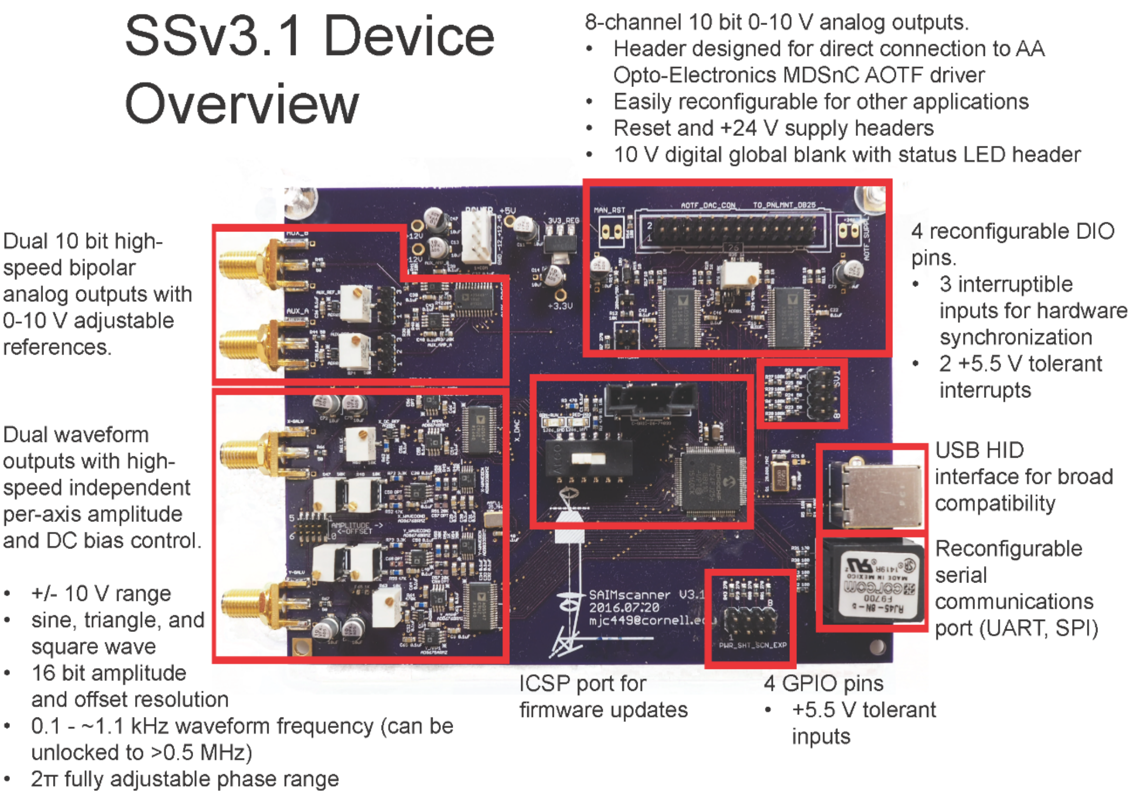


**Fig. S7:** Circuit board layout. All the controller electronics are integrated into a single, compact PCB. The package sizes were chosen to strike a balance between small form factor and serviceability. Hand-building or repairing the electronics is possible with a basic, manual surface-mount device rework station.

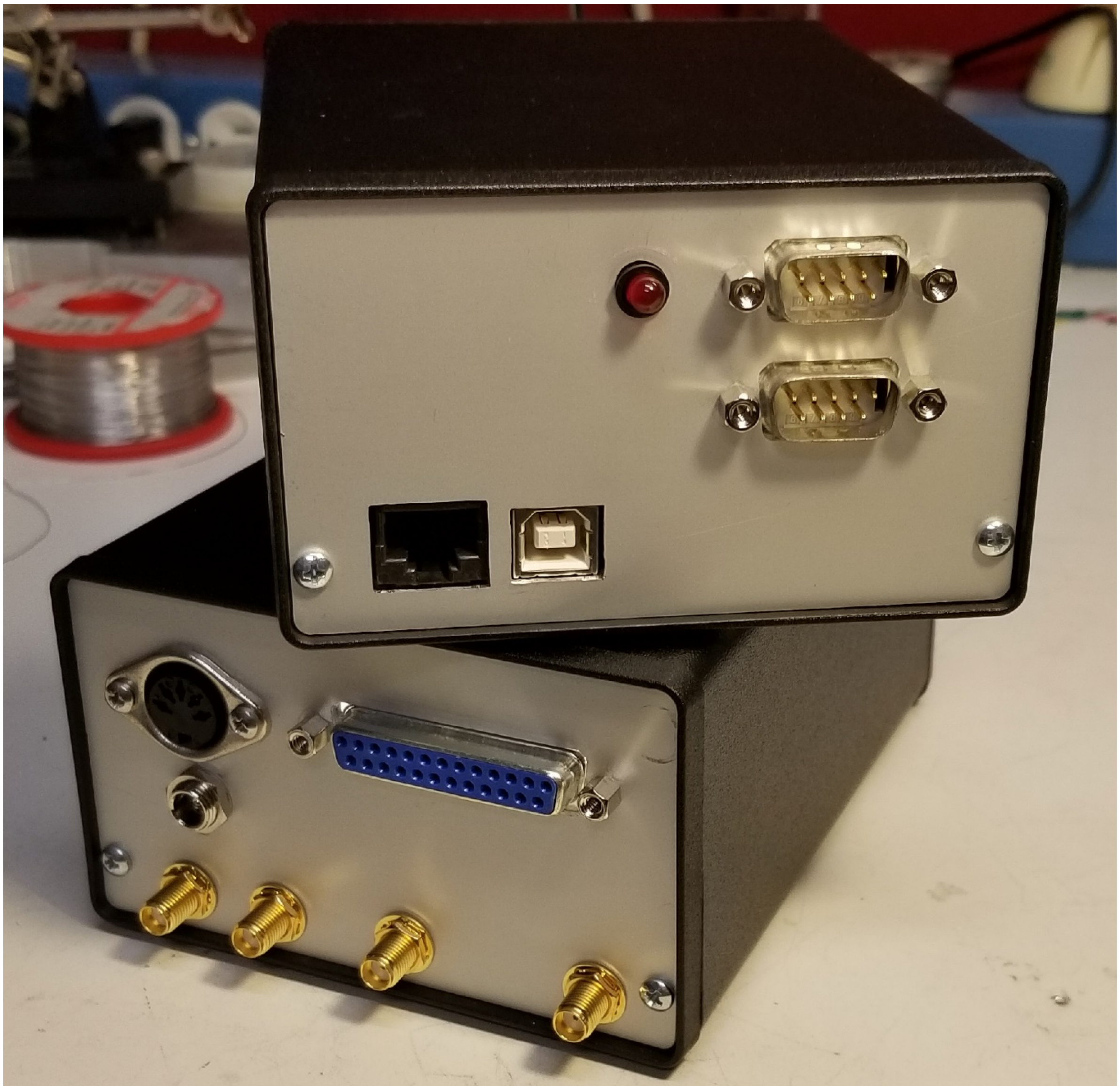

**Fig. S8:** The complete controller assembly with relevant connectors.

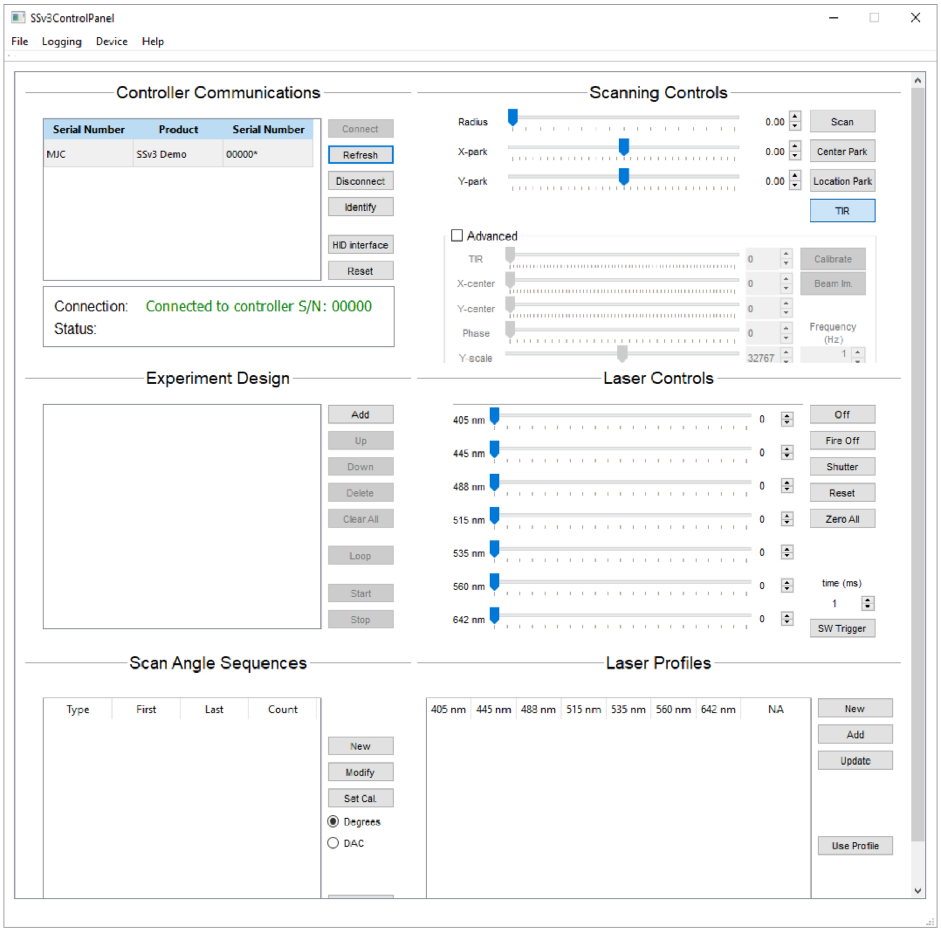


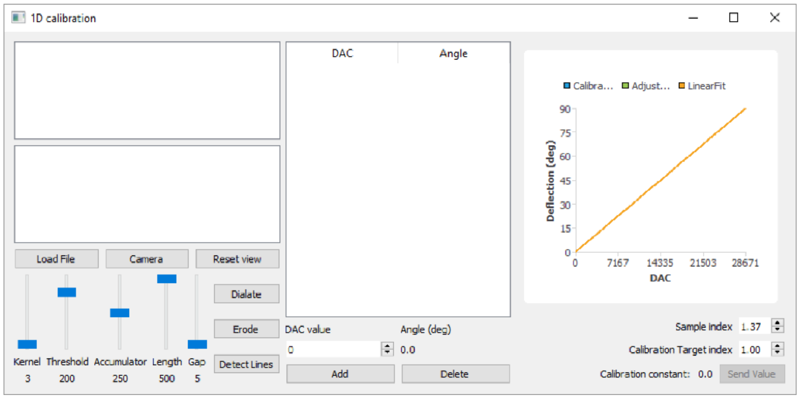


**Fig S9:** Graphical interface for circle scanning applications. Top: the main user interface can be used to create series of scan angles and sets of excitation intensities which can the be used to build complex experimental sequences. The controller can also be run manually through manipulation of the various scan and excitation controls. Bottom: Integrated program and user interface for calibration of the excitation laser beam angle at the sample. Using a webcam pictures of the scanned beam on a vertical surface (i.e. a piece of fluorescent plastic) are taken at various scan amplitudes. The software locates the edges of the scan projection in each image and calculates the calibration constant.
